# A slow and clathrin-dependent mode of compensatory endocytosis in hippocampal boutons at near-physiological temperature

**DOI:** 10.1101/2025.03.31.646354

**Authors:** Martin Kahms, Shrutija Pandey, Jürgen Klingauf

## Abstract

The mode of presynaptic compensatory endocytosis has been controversial for decades. Recently, an ultrastructural study on synaptic membrane retrieval following synaptic vesicle fusion revealed an ultrafast mechanism of endocytosis in mouse hippocampal synapses at physiological temperature. Using live cell imaging of single-vesicle exo- and endocytosis in mouse hippocampal synapses, combined with accelerated optogenetic acidification of synaptic vesicles, we found that compensatory endocytosis occurs mostly through a slow clathrin-dependent and actin-independent mode at both room and physiological temperatures.

## Introduction

To maintain neurotransmission, exocytosed synaptic vesicle (SV) components must be retrieved from the surface by compensatory endocytosis. For decades, clathrin-mediated endocytosis (CME) from the plasma membrane (PM) in a single step has been established as the principal mechanism for this retrieval, first at the frog neuromuscular junction (NMJ) ^1,2^, and later in murine boutons of the central nervous system (CNS) ^3^ and synaptosomes ^4,5^. Electron microscopy (EM) data revealed that SV exo- and endocytosis, though temporally tightly coupled, are spatially segregated: SVs fully collapse and flatten into the PM at the active zone (AZ), and SV proteins and lipids are retrieved elsewhere outside the AZ ^2^. Since clathrin cannot directly bind to the PM or to SV cargo proteins, specific adaptor proteins are required to link clathrin coat formation to the simultaneous selection and concentration of cargo proteins ^6^. A high concentration of cargo is of paramount importance for SV recycling since SVs contain about 40-80 integral membrane proteins with the enormous density of 120 x 10^3^ transmembrane domains per µm^2^ of membrane ^7^. Thus, CME is the only mechanism that enables the high fidelity sorting and concentrating of exocytosed SV proteins to a degree that allows SV retrieval in ‘a single vesicle budding step involving clathrin and dynamin’ ^5^.

More recent results, however, obtained by high-resolution cryo-EM after rapid high-pressure freezing in the nematode *C. elegans* suggest that at room temperature (RT, in the physiological range for *C. elegans*), an ultrafast endocytosis (UFE) mechanism dominates. This process involves dynamin but not clathrin and is completed within 50–100 ms after stimulation, just outside the AZ ^8^. Similar to early EM work of slam-frozen frog NMJs, fusing SVs appeared to fully collapse and flatten within the PM. This concept was later extended to murine hippocampal boutons in culture but only at physiological temperature (PT) and following single AP stimulation ^9^. Thus, it seems that in mammalian synapses at non-physiological temperature and after strong stimulation, slow CME can efficiently and energetically favorably retrieve exocytosed SV components from the PM in a single step. On the other hand, at PT, membrane is retrieved in larger pieces (UFE vesicles have four times the membrane area compared to small SVs) outside the AZ. These vesicles must then fuse with an early endosome, where diluted SV proteins are sorted by adaptors and retrieved through CME ^10^.

The latter clathrin-independent mechanism requires an excess of unsorted endocytosed membrane patches to fuse to endosomes for the reformation of a single fusion-competent SV. Slow CME from these endosomes can retrieve one fusion-competent SV, while the remaining ‘empty’ excess membrane eventually needs to refuse by endosomal exocytosis with the PM to balance membrane and protein trafficking. Hence, UFE is energetically twice as costly compared to CME from the PM, as it requires two dynamin-mediated scission events plus re-fusion of excess membrane. To date no molecular explanation or mechanism has been identified to explain why UFE can proceed in *C. elegans* at low temperatures but not in mammalian synapses, even though proteins involved – such as dynamin – show high sequence homology and retain the same biochemical properties ^11^. Additionally, it remains unknown whether at the frog NMJ, where CME was first described as the mechanism for compensatory endocytosis, at high temperatures (e.g. in a tropical climate or in a hot tub zone), a switch from CME to UFE occurs – and if so, at which exact temperature this binary switch in modes happens. Another significant problem with the UFE model is the intrinsic speed of dynamin-mediated scission. In non-neuronal cells, the buildup of dynamin at the neck of a budding vesicle and subsequent scission has been shown to take on average 10 s ^12^, whereas UFE appears to be completed within 100 ms. To reconcile this, it has been recently suggested that dynamin is pre-assembled at the membrane around the AZ ^13^. However, a detailed molecular model explaining how this preassembled dynamin can reorganize around a neck that only forms after stimulation within 50 to 100 ms and then execute fission has not been developed.

In principle, it should be possible to test the UFE hypothesis using fluorescent exo-endocytosis reporters, such as the pH-sensitive fluorescent protein pHluorin (pHl; ^14^). However, this approach faces several challenges. pHl-based tracers exploit the fact that fusion-competent SVs are acidified by the vacuolar ATPase (v-ATPase), become neutral during fusion pore formation, and are re-acidified following scission. Measurements of re-acidification kinetics, however, have revealed slow time courses in the order of 4 to 7 s ^15,16^. Therefore, in the simplest scenario –where only one mode of retrieval exists such as CME – the reported time constant for the recovery of pHl signals is a convolution of endocytosis, defined by the time point of scission, and the subsequent slow re-acidification. With UFE, the problem of slower reported kinetics is further complicated by the lack of a sorting mechanism in this budding step, which may prevent the retrieval of v-ATPase. As a result, the fluorescence decay in this case would reflect the fusion of vesicles with an endosomal intermediate (pH 6.5), followed by budding from the endosome via CME and final acidification to pH 5.5.

A second problem with pHluorin measurements at the single SV level is the poor signal-to-noise ratio. Most SVs contain only one or two pHl molecules ^17^. With a dynamic range of about 1:20 (at pH 5.5 vs. 7.2) and about 200 SVs in the bouton, and the complement of 10 SVs at the surface (5 % surface pool), we can only expect at best about a 3 % increase in total bouton fluorescence for single SV release, which is typically below the noise level of the baseline signal. However, with high excitation powers single SV exocytosis can be reasonably well measured ^17^. But due to strong photobleaching of surface-resident pHl molecules, endocytosis measurements become significantly confounded. In particular, since many SV contain only one pHl molecule a single step-like bleaching event could be mistaken for a kiss-and-run or UFE event. All published experimental data where high excitation power was used to visualize single SV exocytosis did not avoid photobleaching, and therefore suffered from this issue ^18–21^. Roughly speaking, if individual SV events are observed, pHl molecules at the surface are inevitably bleached. It should be noted that even a for a single endocytic event involving only one retrieved pHl molecule, kiss-and-run or UFE would not manifest itself as a sharp, sub-second fluorescence drop, as often falsely assumed. Instead, the fluorescence would decline with the same slow time course as reacidification, since the protonation-deprotonation reaction of GFP occurs on the tens of microseconds timescale ^22^, and the reacidification of a single SV is not a single stochastic event. Rather, it involves the import (through the vATPase) and simultaneous export (through VGlut in glutamatergic synapses) of more than 1000 protons with transport rates of several 100 per second, given that the stoichiometry of glutamate proton exchange is close to one ^23,24^. Thus with camera integration times of more than 10 ms the expected flickering of the single pHl molecule between protonated and deprotonated states as well as the fast changes in intravesicular proton concentration cannot be resolved. Instead, the measured fluorescence amplitudes reflect the average time the molecule spends in the deprotonated state, which depends on the average proton concentration in that time interval, leading to a gradual fluorescence decline also for single SVs ^25^ containing often only one pHl molecule ^17^.

We thus implemented a two-pronged approach to address the two problems described above, allowing us to faithfully track the fate of pHl molecules post-fusion at both room and near-physiological temperatures. This approach primarily involved measuring single exo- and endocytic events under non-bleaching conditions, where individual events cannot be distinguished from the baseline signal. However, averaging of hundreds of single bouton responses across space and time reliably reveals the average retrieval kinetics. Furthermore, we utilized the optical reporter pHoenix, Synaptophysin1-pHl (Syp1-pHl) fused to the light-activated proton pump Arch3 ^26^. Light activation of pHoenix/Arch3 induces fast acidification of SVs (∼ 1 s) bypassing the intrinsically slow acidification kinetics and bypassing even the need for v-ATPase retrieval along with the pHoenix cargo. With this experimental approach we found that compensatory endocytosis after single action potential stimulation occurs through a single slow mode whose kinetics are best attributable to clathrin-mediated endocytosis at both room and physiological temperatures. Furthermore, we show that under conditions of increased release probability SV cargo sorting is impaired showing. These data indicate that rapid membrane invaginations upon increased amounts of membrane added to the PM by exocytosis do constitute the *bona fide* SV cargo retrieval pathway.

## Results

### Imaging single SV exo- and endocytosis

We first determined the overall kinetics of pHl recovery at RT and PT using Syp1-pHl ^3^ in primary hippocampal neurons with trains of 100 action potentials (APs) at 10 Hz (Fig. 1a). Mono-exponential fits to the decay phases yielded 16.2s for RT and 7.8 s for PT, i.e. a Q_10_ of about 2 (Fig. 1b). We also measured acidification kinetics using the acid quench method ^15^, whereby the surface pool – i.e., the non-endocytosed pool of Syp1-pHl – is rapidly quenched by superfusion with pH 5.5 buffer shortly after stimulation. This reveals the acidification of those SV that had already pinched off at the time of acid application (Fig. 1c and suppl. Fig. 1a). With time constants of 4.2 s for RT and 3.0 s for PT, re-acidification appears to be only mildly temperature-dependent (Fig. 1d).

**Figure 1.**
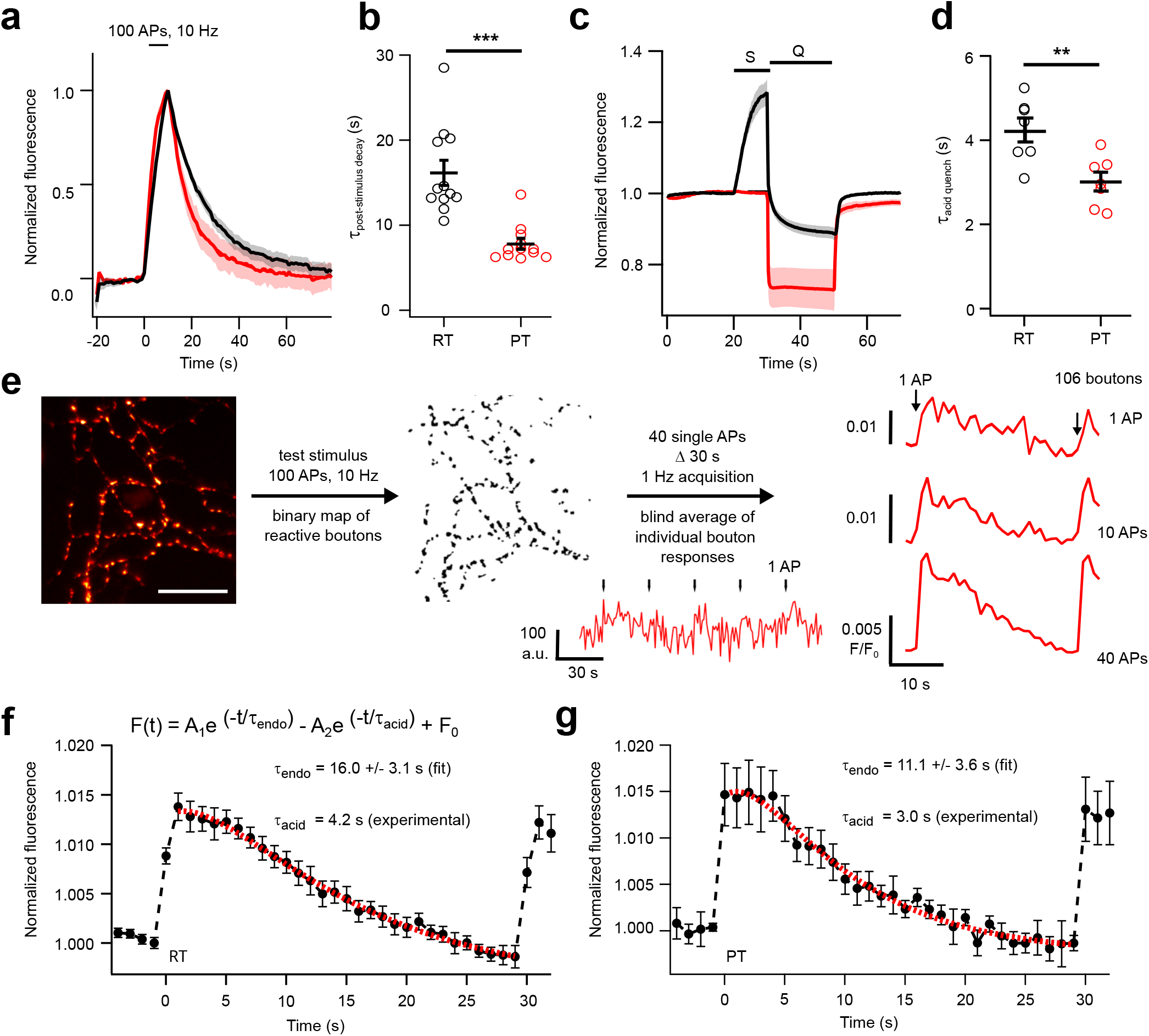
Fluorescence imaging of single vesicle exo- and endocytosis using pHluorin. **a)** Average fluorescence response of Syp1-pHl to a stimulus train of 100 APs at 10 Hz at RT (black; n = 13 coverslips from 3 biological replicates) and at PT (red; n = 12 coverslips from 3 biological replicates). **b)** Time constants of post-stimulus decay were obtained from mono-exponential fits of the recovery phase (τ=16.2 ± 1.5 s at RT and τ = 7.8 ± 0.61 s at PT). *** p <0.005, two-tailed unpaired t-test. **c)** Time course of SV re-acidification determined using the acid quench method at PT. Average fluorescence responses of Syp1-pHl without (red; n=4 coverslips from 2 biological replicates) and with stimulation (black; n=7 coverslips from 2 biological replicates). S denotes field stimulation with 100 APs at 10 Hz, and Q denotes rapid surface quenching immediately after end of stimulation with an impermeant acid at pH 5.5. **d)** Time constants of re-acidification were obtained from mono-exponential fits of the fluorescence decay after end of stimulation and start of the acid quench (τ_acid_ = 4.2 ± 0.2 s at RT and 3.0 ± 0.3 s at PT). ** p <0.05, two-tailed unpaired t-test. **e)** Experimental design for detecting single exo/endocytosis events using Syp1-pHl. Transfected neurons were first stimulated with 100 APs at 10 Hz to obtain kinetic transients of pHl fluorescence as shown in Fig. 1a. A binary mask of reactive boutons was generated from these responses using wavelet transformation (Scale bar: 20 µm). Subsequently, 40 single APs were elicited in a time locked manner at 30 s intervals. Fluorescence signals were recorded at 1 Hz using low illumination intensities and a low duty cycle to prevent photobleaching. Under these conditions, the signal is dominated by shot-noise. Fluorescence responses to single APs from the identified reactive boutons were blindly averaged, resulting in a signal-to-noise ratio increase proportional to the square root of n, where n is the number of individual bouton responses. **f)** Ensemble-averaged fluorescence trace of neurons expressing Syp1-pHl in response to single APs at RT (n=7 coverslips from 2 biological replicates). A double-exponential fit to the decay curve (red, shown as fit of the average curve) models SV endocytosis and re-acidification as two sequential first-order processes. τ_acid_ was fixed to 4.2 s during the fitting (experimental results shown in 1d) to determine τ_endo_. Fitting individual curves yielded τ_endo_ = 16.0 ± 3.1 s. **g)** Ensemble-averaged fluorescence trace of Syp1-pHl responses to single APs at PT (n=6 coverslips from 2 biological replicates). Double-exponential fitting of the individual curves with τ_acid_ fixed to 3 s resulted in τ_endo_ = 11.1 ± 3.6 s All data are shown as mean ± s.e.m.

Next, we determined the recovery kinetics for single APs. We first localized responsive boutons by a applying a burst of 100 APs at 10 Hz, either at RT or PT, and then recorded Syp1-pHl responses to 40 single APs, administered every 30 s under low illumination intensity (2 W/cm^2^) with a duty cycle of 0.25%. Under these conditions, no photobleaching of the surface pool was observed (Fig. 1e). Averaging at RT yielded an average single AP response with kinetics similar to those measured for Syp1-pHl in response to 100 APs at 10 Hz (Fig. 1f), showing that at RT the slow endocytic rate dominates the signal over the re-acidification. Responses to single APs, however, reveal a marked plateau, especially at PT after the exocytic increase (Fig. 1g). This is expected and should reflect the slow re-acidification of the internalized SVs ^15^. The measured decay kinetics of Syp1-pHl represent a convolution of SV endocytosis followed by re-acidification in case of CME or, alternatively in the case of UFE, the slow transport of Syp1-pHl molecules from the PM to SVs through an endosomal intermediate (pH 6.5) accompanied by slow acidification. Since exocytosis after a single AP occurs within ms, i.e. 2 to 3 orders of magnitude faster than UFE or CME, the pHl transient can be considered as the response to an impulse function. Thus, the convolution simplifies to a double-exponential decay for two sequential first-order reactions representing endocytosis and re-acidification, provided SV recycling proceeds in a single budding step by a single endocytosis mechanism. Therefore, a double exponential fit should yield both the time constant of re-acidification as well as the time constant of endocytosis. We used the above measured re-acidification time constants for the faster exponential and obtained endocytosis time constants of 16 s for RT and 11 s for PT by least-square fitting (Fig. 1f,g). While the time constant at RT is identical to the one obtained for 100 APs, endocytosis in response to single APs appears to be somewhat slower than for 100 APs at PT. Such use-dependent acceleration of endocytosis for stronger stimulation has been described ^27^. To test the robustness of our experimental approach, we averaged the data set shown in Fig 1e with Δ_average_ of 31 s instead of 30 s and found that the resulting trace was dominated by noise (suppl. Fig. 1b). Furthermore, we repeated the experiment in the presence of the v-ATPase blocker bafilomycin (Baf) and observed the anticipated step-like profile (suppl. Fig. 1c).

### Imaging single SV exo- and endocytosis under accelerated acidification conditions

Next, we carried out analogous measurements using pHoenix. Activation of the Arch3 proton pump with 561 nm light leads to additional proton pump activity, which helps to better deconvolve acidification and endocytosis (Fig. 2a). We first determined the SV acidification kinetics of pHoenix. Optogenetic acidification under acute block of the v-ATPase with Baf yields more complex acidification kinetics that are no longer are mono-exponential. However, a double-exponential gives a very good fit, judging from the residuals (Fig. 2b), with a rapid component (τ_fast_ ∼ 1 s) being the dominant one (70 % amplitude), followed by a slower component (τ_slow_ ∼ 8 s), which accounts for only 30 % of the amplitude at PT. Similar acidification kinetics for pHoenix were found at RT (suppl. Fig 2a). Thus, we found pHoenix acidifies SVs three times faster compared to the endogenous v-ATPase (suppl. Fig. 2b), fully in line with data reported earlier ^26^. If anything, re-acidification must be even faster without the Baf block of the endogenous v-ATPase. Without Arch3 activation, pHluorin signals of pHoenix look similar in response to 100 APs at 10 Hz compared to Syp1-pHl (Fig. 2c). Mono-exponential fits to the decay phases yield 16.9 s for RT and 9.1 s for PT (Fig. 1d), not significantly different to the values obtained for Syp1-pHl. When these experiments were performed under constant activation of Arch3 by 561 nm light, a slight but not significant decrease in τ_post-stimulus decay_ was observed, indicating that endocytosis is the main rate-limiting step for this burst of APs (Fig. 2d and suppl. Fig. 1c).

**Figure 2.**
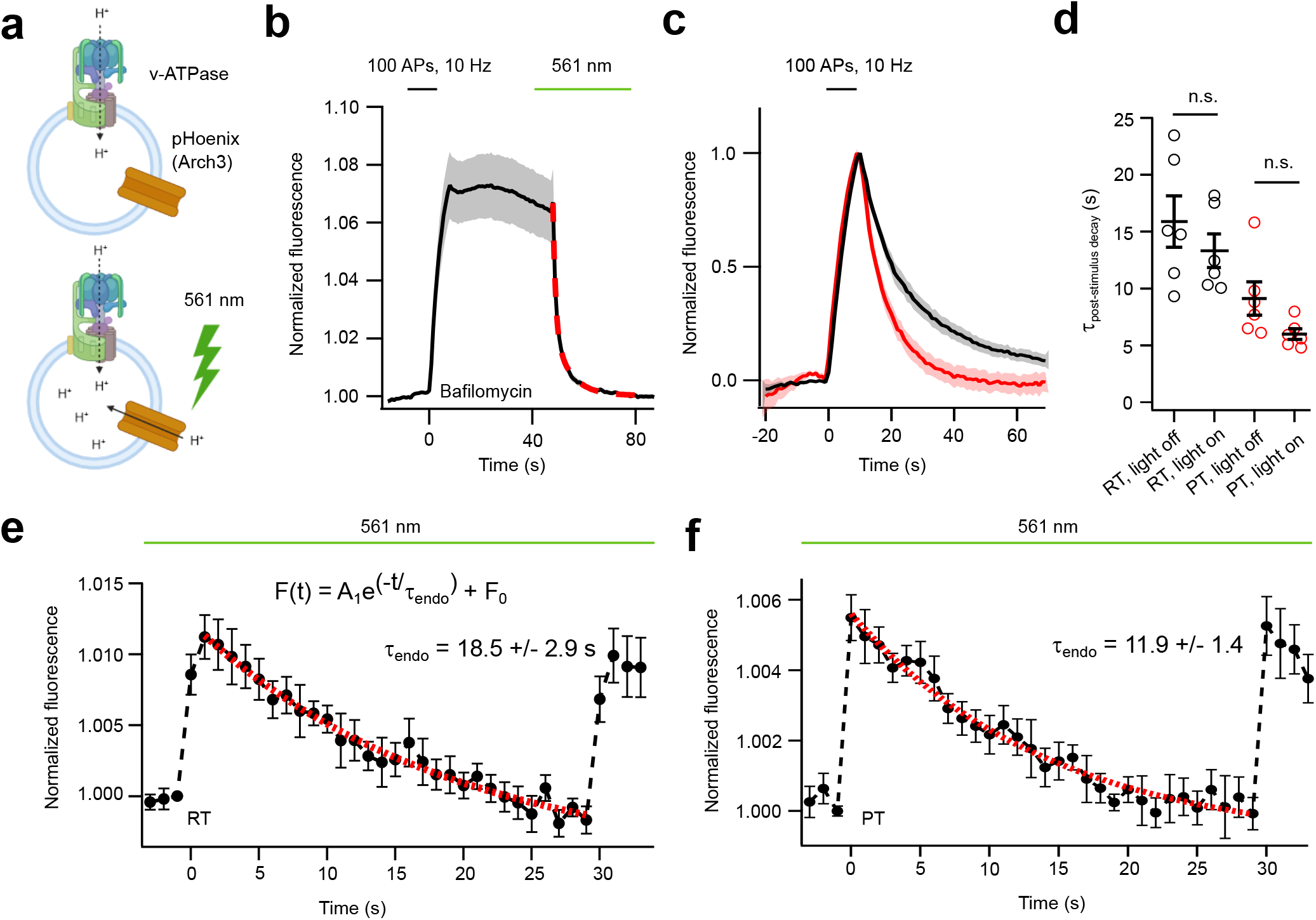
Imaging single vesicle exo-endocytosis with optogenetically accelerated synaptic vesicle acidification. **a)** Schematic illustrating the optogenetic activation of pHoenix. The proton pump Arch3 is targeted to SVs by fusion with Syp1-pHl. Upon activation of pHoenix/Arch3 with 561 nm light, its proton pump activity, in addition to the endogenous v-ATPase, results in faster SV acidification. Created with BioRender.com. (**b)** Acidification kinetics of the optogenetic proton pump pHoenix/Arch3 at PT. Hippocampal neurons were stimulated with 100 APs at 10 Hz in presence of 1 µM Baf. After completion of endocytosis, the proton pump Arch3 was activated by 561 nm light to estimate the optogenetically induced acidification rate of pHoenix (n=10 coverslips from 3 biological replicates; red: double exponential fit of the average curve). Fitting the individual curves to a double-exponential function yields τ_fast_ = 1.3 ± 0.3 s (66 ± 9 % of amplitude) and τ_slow_ = 8.4 ± 1.1 s (34 ± 9 % of amplitude). **c)** Average fluorescence response of pHoenix to a stimulus train of 100 APs at 10 Hz at RT (black; n = 6 coverslips from 2 biological replicates) and at PT (red; n = 12 coverslips from 3 biological replicates) without Arch3 activation. **d)** Endocytosis time constants were obtained from mono-exponential fits of the recovery phase for measurements without and with Arch3 activation (τ_endo, light off_ =15.9 ± 2.3 s at RT and τ_endo, light off_ = 9.1 ± 1.5 s at PT; τ_endo, light on_ =13.3 ± 1.5 s at RT and τ_endo, light on_ = 5.9 ± 0.5 s at PT). n.s.: not significant, two-tailed unpaired t-test. **e)** Ensemble-averaged fluorescence trace of neurons expressing pHoenix in response to single APs at RT. The averaged trace shows the fluorescence increase in response to single APs followed by subsequent decay of fluorescence due to SV endocytosis and re-acidification by Arch3 (continuous activation) and the endogenous v-ATPase. A mono-exponential fit (red, shown as fit of the average curve) to the individual decay curves yields τ_endo_ = 18.5 ± 2.9 s at RT (n=7 coverslips from 3 biological replicates). **f)** Same measurements as in e) but at PT. A mono-exponential fit (red, shown as fit of the average curve) to the individual decay curves yields τ_endo_ = 11.9 ± 1.4 s at RT (n=9 coverslips from 3 biological replicates). All data are shown as mean ± s.e.m.

If our interpretation of the measured double-exponential decay for single AP measurements with Syp1-pHl was correct, rapid optogenetic re-acidification by pHoenix should affect only the plateau phase in single AP measurements, but not the late phase. In fact, the plateau phase should disappear, resulting in a mono-exponential decay reflecting the probability density function for the times of scission, provided one endocytosis mechanism is dominant. In other words, if UFE was the dominant form of compensatory endocytosis for single AP stimulation at PT, repeating the experiment in Fig. 1g with light-activated Arch3 acidifying the endocytosed SVs along with the v-ATPase should result in an immediate drop of the pHl response to about 20 % with a time constant of 1 s upon a single AP. However, we observe a slow mono-exponential decay with time constants of 18.5 s for RT and 11.8 s for PT respectively, nearly identical to those obtained for the slow component in the absence of Arch3 activation (suppl. Fig. 2c,d) and the ones we measured for Syp1-pHl (Fig. 1f,g). Remarkably, the plateau phase is now lacking, as predicted by the double-exponential model of re-acidification and endocytosis, proving much faster acidification kinetics during light activation of Arch3. To ensure that we did not miss a rapid drop in amplitude at a recording frequency of 1 Hz, we repeated the measurements at PT under Arch3 activation at 10 Hz, collecting only 40 frames around the stimulus to maintain the duty cycle (suppl. Fig 2e). In this case, only 4 APs were elicited with 30 s intervals. The slow decay observed in the first 3 s matches well the decay observed at 1 Hz (Fig. 2f) and that we did not miss an immediate drop in amplitude when sampling at 1 Hz. If pHoenix was retrieved solely by UFE, the expected fluorescence decay time course would correspond to the convolution of the measured reacidification time course at PT in presence of Baf (Fig. 2b) and the UFE time course (Fig. 3a). The expected fluorescence decay should be even faster, since the vATPase was not blocked in these experiments. Given the noise level in our measurements, we conclude that in murine hippocampal boutons at PT, UFE does account for less than 15 % of compensatory endocytosis in response to single APs, if at all.

**Figure 3.**
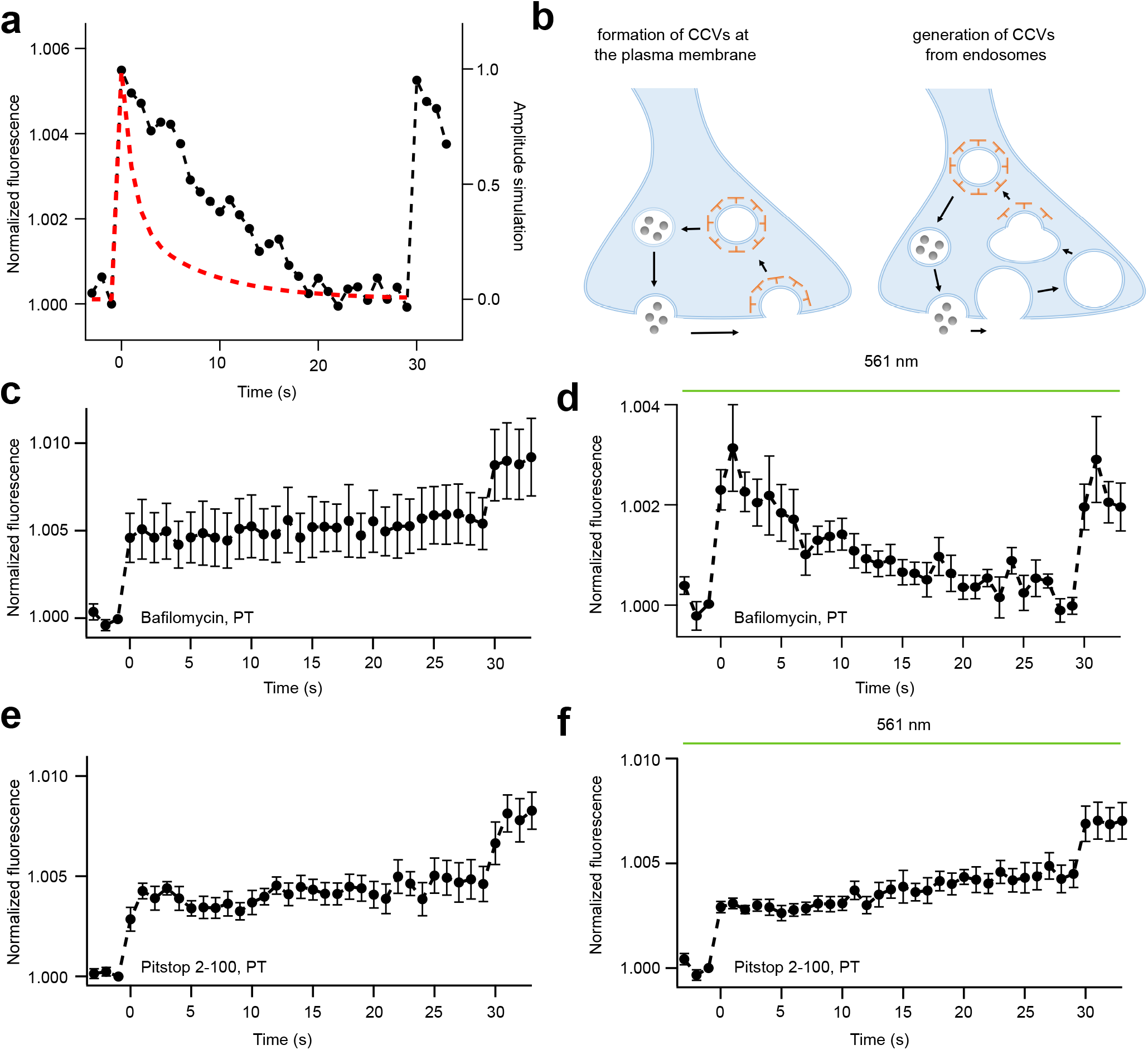
Simulation of UFE and role of clathrin. **a)** Simulated time course of pHoenix fluorescence for single APs and optogenetic acidification under the assumption of ultrafast endocytosis only (red, convolution of pHoenix acidification and ultrafast endocytosis kinetics). For comparison, the measured single AP time course at PT with light-induced pHoenix activation shown in Fig. 2f is plotted (black). **b)** Two scenarios illustrating the different action of clathrin assumed for different endocytosis modes. The classical clathrin mediated endocytosis pathway involves clathrin (blue) operating at the PM. The ultrafast endocytosis mechanism is followed by clathrin mediated budding from endosomes. Ensemble-averaged fluorescence traces of neurons expressing pHoenix in response to single APs at PT in presence of 1 µM Bafilomycin shown in **c)** without light activation of pHoenix and in **e)** with light activation of pHoenix. The plateau step-like fluorescence profile observed without light activation of pHoenix can be converted into a fluorescence decay upon light delivery, indicating that the pHoenix cargo has been endocytosed and is located in acidifiable SVs Ensemble-averaged fluorescence traces of neurons expressing pHoenix in response to single APs at PT in presence of 45 µM Pitstop 2-100 shown in **e)** without light activation of pHoenix and in **f)** with light activation of pHoenix. In both cases, a step-like profile is observed, indicating that the pHoenix cargo remains at the PM and is not located in acidifiable endosomal structures. Ensemble-averaged fluorescence traces of neurons expressing pHoenix in response to single APs at PT in presence of 1 µM Bafilomycin shown in **e)** without light activation of pHoenix and in **f)** with light activation of Phoenix. The plateau step-like fluorescence profile observed without light activation of pHoenix can be converted into a fluorescence decay upon light delivery, indicating that the pHoenix cargo has been endocytosed and is located in acidifiable SVs (n=7-8 coverslips each from 2 biological replicates). All data are shown as mean ± s.e.m.

### Block of clathrin function inhibits SV scission at the plasma membrane

Recycling of SVs by UFE involves clathrin, but only at a later step after scission from the PM – specifically, during the budding of SVs from endosomes ^10^ (Fig. 3b). Therefore, we next sought to investigate the role of clathrin during SV endocytosis at PT. As a first control experiment, we conducted measurements in the presence of the v-ATPase blocker Baf (1 µM) at PT, resulting in a step-like profile (Fig. 3c). Reacidification was fully restored when Arch3 was activated by light (Fig. 3d). This result indicates that endocytosis is normal in the presence of Baf ^28,29^ and that the pHoenix cargo has been sorted correctly to SVs. Next, we repeated these experiments using the clathrin inhibitor Pitstop 2-100 ^30^ (Fig. 3e). Pitstop 2-100 reliably blocked SV endocytosis at a concentration of 45 µM at RT for a bout of APs (100 APs, 5 Hz, suppl. Fig. 3a,b). Without Arch3 activation, like under Baf, we observed a step-like fluorescence profile indicating that blocking clathrin function indeed impairs SV reformation (Fig. 3e). However, when these measurements were performed under constant Arch3 activation, reacidification could not be restored (Fig. 3d). Thus, when impairing clathrin function, pHoenix could not be retrieved from the PM. It is however unlikely that in these experiments exocytosis did not trigger compensatory endocytosis. Rather exocytosed excess membrane was taken up by a clathrin-independent mode of endocytosis without cargo sorting capability like UFE. Overall, our results show that compensatory endocytosis of SVs in response to a single stimulus follows kinetics best attributed to a single, slow, and clathrin-dependent mode of endocytosis at RT and PT.

### High release probability triggers incomplete cargo retrieval

The rapid formation of uncoated pits with four-fold membrane area compared to SVs has been already observed in early studies at the frog NMJ ^2^, where almost a third of exocytosed membrane was retrieved by this fast mechanism. However, unlike clathrin-coated pits, these pits seen in freeze-fracture replicas did not enrich cargo and were thought to spontaneously form due to massive release under unphysiological stimulation (in presence of the K^+^ channel blocker 4-AP) to compensate for the imbalance of outer and inner leaflet lipids added to the PM. In line, simulations on UFE showed that two or more SVs have to synchronously fuse to create enough lateral compression to trigger UFE ^31^, while under physiological conditions CNS synapses release probability is low with typically only one or no SV fusing ^32^. We therefore sought to increase the release probability by elevating external Ca^2+^, thereby potentially eliciting UFE. When repeating the experiment in Fig. 2f in presence of 8 mM external Ca^2+^, we observed incomplete cargo retrieval indicating that a large fraction of exocytosed membrane was retrieved in a compensatory way without cargo (Fig. 4a). However, when we disrupted presynaptic actin organization by application of 10 µM Latrunculin A (Lat), a treatment which has been shown to prevent initiation of UFE ^31^, cargo retrieval becomes compensatory again even at 8 mM external Ca^2+^ (Fig.4b). The fluorescence decay can again be well described by a single-exponential with a time constant of 6.7 s, reflecting the acceleration of CME upon increased Ca^2+ 33^. Interestingly, however, a slight plateau in the first second or so after exocytosis is observed (Fig.4a), absent in the trace in 2 mM Ca^2+^ (Fig. 2f) as well as after application of Lat (Fig. 4b). This indicates that during this time membrane is retrieved by a different rapid mechanism lacking cargo-sorting capability and that only at later times CME ensues until the membrane tension is fully restored, therefore leaving quite some cargo behind. This supports the notion, that under non-physiological conditions or in general under conditions of very high release probability UFE is triggered, which however fails to enrich SV cargo.

**Figure 4.**
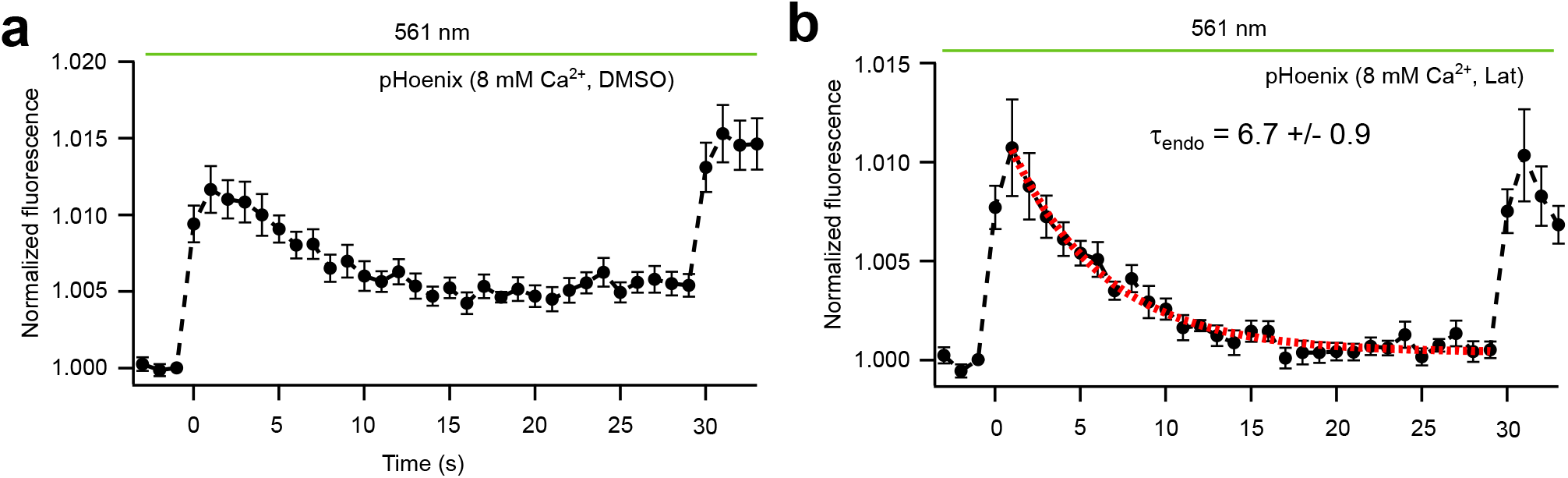
UFE endocytosis and membrane tension. **a)** Ensemble-averaged fluorescence traces of neurons expressing pHoenix in response to single APs at PT in presence of 8 mM external Ca^2+^and under light activation of the pHoenix pump. The incomplete post-stimulus fluorescence decay indicates only partial cargo retrieval. **b)** Ensemble-averaged fluorescence pHoenix traces in response to single APs at PT in presence of 8 mM external Ca^2+^ and 10 µM Latrunculin A, and under light activation of the pHoenix pump. Full post-stimulus fluorescence decay can be restored indicating complete cargo retrieval (n=9-10 coverslips each from 2-3 biological replicates). A mono-exponential fit (red, shown as fit of the average curve) to the individual decay curves yields τ_endo_ = 6.7 ± 0.9 s All data are shown as mean ± s.e.m.

## Discussion

The maintenance of the complete SV proteome is critical to maintain synaptic transmission. Therefore, it has long been postulated that only CME can ensure the proper sorting of multiple vesicle cargo proteins ^3,34,35^. But an early study at the frog NMJ already reported that, in addition to slow CME, fast clathrin-free invaginations could also be observed ^2^. These invaginations, however, appeared to be predominantly free of cargo. Consequently, it was postulated early on that these fast membrane invaginations were merely an ‘abnormal manifestation’ resulting from the massive incorporation of SV membrane into the PM (these experiments were performed in the presence of the potassium-channel blocker 4-AP). Decades later, flash and freeze experiments in conjunction with optogenetic stimulation confirmed such rapid membrane invaginations 50-100 ms after exocytosis in CNS synapses. This type of ultrafast endocytosis was defined as a *bona fide* SV retrieval pathway after single AP stimulation at PT, in contrast to CME which operates only at RT ^9^. Generally speaking, it is evident that such rapid invaginations exist, as capacitance measurements also revealed a fast component of membrane retrieval at PT ^36^. Moreover, the subsequent fate of these rapidly invaginated and pinched off structures, in terms of endosomal recycling, is certainly plausible ^10^. However, the key question remains: does this pathway represent the *bona fide* SV cargo recycling pathway. To answer this, it is essential to switch from a membrane-centered to a cargo-centered perspective. In this context, experiments using pHluorin tagged SV molecules have been performed. Some of these studies indeed reported an ultrafast cargo retrieval after single AP stimulation. However, these data typically show an instantaneous pHluorin fluorescence drop for ultrafast events in less than 100-200 ms ^18,21^. Since pHluorin reports reacidification, these values are inconsistent with the acidification kinetics of v-ATPase ^15,16^ as well as with the fast proton dynamics of GFP-based molecules ^22^. Therefore, it is highly likely that these measurements are confounded by photobleaching as the time course of SV acidification and the associated gradual decay of pHluorin fluorescence in the seconds range has been demonstrated in single isolated SVs of transgenic mice expressing pHluorin in the vesicular lumen ^25^. When we measured single SV retrieval kinetics for the most abundant SV cargo, Synaptophysin1 (fused to pHluorin), under non-bleaching conditions, our data at both RT and PT are most compatible with two sequential first or processes describing acidification as well as slow endocytosis. Furthermore, additional experiments with the optogenetic proton pump pHoenix showed no evidence of an ultrafast pathway that also carries significant cargo. Instead, our findings indicate that also at PT clathrin, which is involved in cargo recycling, is primarily active at the PM rather than in a later step at endosomes. Notably, the single SV retrieval kinetics observed for pHoenix under accelerated acidification condition closely match the distribution of scission waiting times measured by total internal reflection fluorescence microscopy (TIRFM) for endogenously tagged Dynamin I-EGFP at PT in the purely presynaptic TIRFM-amenable Xenapse preparation ^37,38^. Of course, we cannot entirely rule out the existence of a small fraction of ultra-fast membrane retrieval component in parallel under our measuring conditions. However, our data suggest that this is not the primary recycling route for SV cargo following a single stimulus, neither at RT nor at PT, because the slow mode seen in our data is fully compensatory with respect to cargo retrieval, implying that also the bouton membrane area is restored. Additional UFE would thus cause shrinkage of the bouton surface, which is not plausible. In line, simulations and experiments have revealed that membrane compression upon exocytosis of multiple SVs triggers ultrafast endocytosis and that a stable actin cortex is required to induce ultrafast membrane retrieval ^31^. Initial experiments using high-resolution cryo-EM after rapid high-pressure freezing utilized optogenetic stimulation in presence of elevated extracellular Ca^2+ 9^. And already here it was observed that the fraction of UFE scales with the amount of membrane added by exocytosis. However, simulations predicted that two or more SVs have to fuse to trigger ultrafast endocytosis ^31^. Thus, the crucial point in interpreting the biological relevance of UFE is determining the number of primed SVs that fuse under physiological conditions in response to a single AP. *Zap and freeze* experiments (electrical single AP stimulation followed by rapid high freezing) revealed release of 1-2 SVs under physiological Ca^2+^ concentrations per single active zone during single AP delivery ^39^. Likewise electrophysiological recordings at single glutamatergic synapses at PT reveal release of only 1-2 SVs upon single APs ^32^. In our experiment, a slightly elevated Ca^2+^ concentration of 2 mM does not appear to induce multiple release events to an extent that significantly triggers UFE. However, at strongly elevated Ca^2+^ concentrations, UFE is induced to rapidly compensate for the excess vesicular membrane added to the PM albeit at the expense of cargo sorting. Thus, experimental strategies relying on high release probability (elevated Ca^2+^, optogenetic stimulation) risk overestimating the proportion of UFE that occurs under physiological conditions. Furthermore, the finding that UFE is not the *bona fide* cargo retrieval route underlines that it is important to switch from a membrane-centered to a cargo-centered perspective.

## Methods

### Plasmids

pLenti-Synapsin-pHoenix-WPRE was a gift from C. Rosenmund (Addgene plasmid #70111). pEGFP-Synaptophysin1-pHluorin was a gift from L. Lagnado (Addgene plasmid #24478). Lentiviral packaging and envelope vectors pCMVΔR8.2 and pMD2.G were a gift from Didier Trono (Addgene plasmid # 12263 and plasmid #12259). Large-scale DNA purification was performed using an endotoxin-free plasmid purification kit (Qiagen).

### Mice

Breeding of C57BL/6N mice was conducted at the Central Animal Experimental Facility of the University Hospital Muenster according to the German Animal Welfare guidelines (permit number AZ81-02.05.50.20.019, issued by ‘Landesamt für Natur, Umwelt und Verbraucherschutz NRW’, Duesseldorf, Germany). Both male and female mice were used for primary neuronal cultures.

### Primary hippocampal neurons

High-density dissociated cultures of mouse hippocampal neurons were prepared from CA3/CA1 regions of 1 to 3-day-old mice by trypsin digestion. Neurons were plated on Matrigel (Corning)-coated coverslips in Minimal essential medium (MEM, Gibco) supplemented with 5 g/l glucose, 200 mg/l NaHCO_3_, 100 mg/l transferrin (Calbiochem), 25 mg/l insulin (Sigma), 10% fetal bovine serum (Sigma-Aldrich), and 2 mM L-glutamine (Sigma). Subsequently, the medium was gradually exchanged for Neurobasal-A medium (Gibco) supplemented with 2% B27 (Gibco), 2 mM Glutamax (Gibco), and 0.1% penicillin/streptomycin (Gibco). On day 3 *in vitro* (DIV), 2 µM cytosine arabinoside (Sigma) was added to inhibit glial cell proliferation.

### Transfection of hippocampal neurons

Transfection was performed on DIV 4 by calcium phosphate precipitation. To this end, neurons were preincubated in pure Neurobasal-A medium without supplements. A precipitate was formed by mixing 3 µg of plasmid DNA, 2.5 µl 2.5 M CaCl_2_, and 19.5 µl dH_2_O per each well of a 12-well plate. Then, an equal volume of 2x BBS solution (50 mM BES, 280 mM NaCl, 1.5 mM Na_2_HPO_4_, pH 7.0) was added dropwise while gently vortexing. The precipitate was allowed to form for 20 min. The complexes were mixed with 450 µl of Neurobasal-A medium per well and 500 µl of the solution was added to each well. Plates were returned to the incubator (5% CO_2_, 37 °C) for 20 minutes and washed three times with prewarmed HBSS solution (w/o Ca^2+^ and Mg^2+^). Finally, the cells were transferred back into their previously saved conditioned growth medium. Neurons were imaged at DIV 15–25.

### Viral production

HEK293FT cells were transfected at approximately 70 % confluency on 10 cm round culture dishes using TransIT-293 transfection reagent (Mirus). After 72 hours, the medium was collected from 4-6 plates and filtered through a 0.4 µm filter. Afterwards, the medium was centrifuged at 25.000 rpm for 90 minutes in an ultracentrifuge (Beckman) using a fixed angle rotor type 45I (Beckman). The supernatant was removed and the pellet was resuspended in 100-200 µl Dulbecco’s phosphate-buffered saline (DPBS, Gibco). Aliquots were stored at -80°C until use.

### Viral transduction of neurons

Transduction of neurons was performed on DIV 1-2. Coverslips were transferred into a plate containing 1 ml Neurobasal-A medium supplemented with 2% B27 and 2 mM Glutamax (growth medium) per well. 1-3 µl of virus solution was added, and cells were incubated in a humidified incubator at 37 °C and 5% CO_2_ for 5 hours. Then, cells were washed 3 times with growth medium and put back into their previously saved conditioned growth medium.

### Epifluorescence microscopy

All experiments, unless otherwise stated, were carried out in modified Tyrode’s solution (25 mM HEPES, 119 mM NaCl, 2.4 mM KCl, 2 mM CaCl_2_, 2 mM MgCl_2_, 30 mM glucose, pH 7.4). Experiments were performed at either RT (21 ± 1 °C) or PT (35 ± 1 °C) within a heating chamber equipped with temperature sensor. Neurons were stimulated by electric field stimulation (platinum electrodes, 10 mm spacing, 1 ms pulses of 50 mA, alternating polarity at 20 Hz) applied by a constant current stimulus isolator (WPI A 385, World Precision Instruments). 10 μM 6-cyano-7-nitroquinoxaline-2,3-dione (CNQX, Tocris Bioscience) and 50 μM D,L-2-amino-5-phosphonovaleric acid (AP5, Tocris Bioscience) were added to prevent recurrent activity. Solution exchange for acid-quench experiment was achieved through a three-barrel glass tubing perfusion system controlled by a piezo-controlled stepper device (SF778, Warner Instruments). Acidic solution (pH 5.5) was prepared by substituting HEPES with 2-(N-morpholino) ethane sulfonic acid (MES). Imaging was performed on an inverted Nikon Ti Eclipse microscope equipped with a fiber-coupled laser box (Omicron). Fluorescence emission was detected using a 60x water immersion objective (Nikon Apo, NA 1.2), a quadband filter set (AHF Analysentechnik) together with either a sCMOS camera (ORCA Flash, Hamamatsu) or an EMCCD camera (Evolve, Photometrics, gain set to 600) controlled by micromanager software with a digital pixel size of 217 x 217 nm^2^ (2x2 binning mode for the sCMOS camera). An optosplit (Cairn Research) with an additional dichroic beamsplitter (570 nm) and emission filters (pHluorin channel (525/45 nm) and mKate channel (612/69 nm)) was used in case of pHoenix measurements. For image acquisition using bouts of stimulation, pHluorin and pHoenix were excited with a 488 nm laser (Phoxx, Omicron), and Arch3 was photoactivated by excitation with a 561 nm laser (Cobolt Jive, 1.1 W/cm^2^). Images were acquired at 1 Hz with an exposure time of 25 ms. For acid-quench experiments, images were acquired at 5 Hz, with an exposure time of 25 ms. For experiments monitoring pHluorin and pHoenix responses to single APs, images were acquired at 1 Hz with an integration time of 25 ms per frame. For each coverslip two data sets were aquired, the first one consisting of a 50-100 APs at 10 Hz stimulus (100 images) and the second consisting of 40 cycles of single AP stimulation with an interval of 30 s between individual APs (1200 images). Single APs were delivered in a time-locked manner 5 ms before the first post-stimulus frame was captured for 25 ms. Image acquisition under no bleaching conditions (short duty cycle of 25 ms/s combined with imaging at 1 Hz and low illumination intensities of 2 W/cm^2^) was complemented with hardware synchronization using the Arduino Uno microcontroller unit (Arduino IDE 1.6.0, Arduino, www.arduino.cc). When imaging under such low light conditions the photon shot noise becomes dominant and we applied an averaging approach to increase the signal-to-noise ratio (SNR). The photon shot noise is Poisson distributed with a square root relationship between signal and noise. Thus, averaging a sufficiently high number of exo-endocytic events reduces the photon shot noise while increasing the signal.

### Image and Data analysis

Drift correction was performed in Fiji/ImageJ (https://fiji.sc/) using the StackReg or TemplateMatching plugin. Quantitative analysis of pHluorin and pHoenix experiments was performed with custom-written macros in Igor Pro (Wavemetrics) as described previously ^40^. Functional boutons were defined as regions in which the fluorescence intensity changes during stimulation. The difference image, created by subtraction of an average of 5 images before and after stimulation was subject to an á-trous wavelet transformation ^41^. In the resulting image mask, spots represent putative functional boutons. Only spots with areas between 4 and 20 pixels were accepted for further analysis. The mask was then overlapped with time lapse images to extract single bouton fluorescence transients. All calculated time courses were visually inspected for correspondence to individual functional boutons. Only experiments with more than 50 active boutons were considered for analysis. For analysis of single AP measurements, a mask of responding boutons (from 50-100 APs 10 Hz stimulus) was blindly used to analyze fluorescence intensities in response to single APs. An averaging approach with a delta of 30 s (interval between single APs) was applied in a time-locked manner to increase the signal to noise ratio (SNR), wherein multiple cycles of single APs (30 s interval between each single APs) were averaged for each measurement.

### Quantification and statistical analysis

Data summation and statistical analyses were performed using Igor Pro (Wavemetrics). Data are shown as mean values ± SEM. The number of total coverslips is denoted as n and in addition, the number of individual cultures the coverslips originate from is given (biological replicates). All data passed statistically analyzed passed the normality test (Kolmogorov-Smirnov) and statistical significance was evaluated using a two-tailed unpaired Student’s test at p < 0.05. n.s. denotes non-significance.

## Figure legends

**Suppl. Figure 1.**
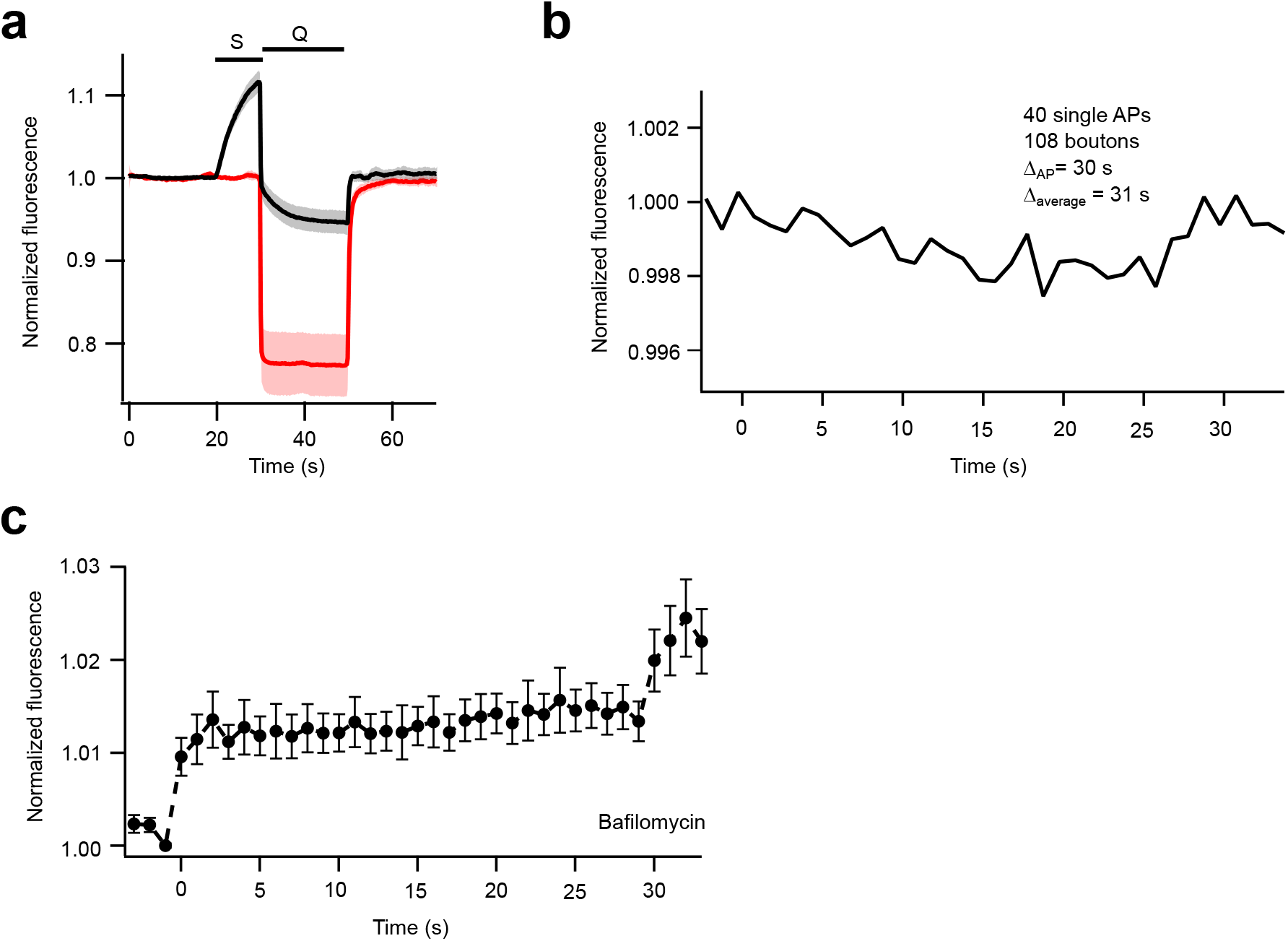
Synaptic vesicle acidification kinetic at room temperature and validation of single vesicle imaging approach. **a)** Time course of SV re-acidification measured using the acid quench method at RT. Average fluorescence responses of Syp1-pHl without (red; n=4 coverslips from 2 biological replicates) and with stimulation (black; n=7 coverslips from 2 biological replicates). S denotes field stimulation with 100 APs at 10 Hz, and Q denotes rapid surface quench immediately after end of stimulation with an impermeant acid at pH 5.5. **b)** Resampling and averaging of data shown in Fig.1d with a Δt other than 30 s (Δt of single AP delivery) results in only noise (shown for a Δt of 31 s). Note that only averaging in a time-locked manner results in an increased signal-to-noise ratio, which finally enables detection of single exo-endocytosis events. **c)** Average fluorescence traces for Syp1-pHl in response to single APs in presence of Baf at RT. Repeated single AP stimulation with an interval of 30 s averaging of single bouton responses results in a staircase-like increase in fluorescence due to block of re-acidification by block of v-ATPase function (n=7 coverslips from 3 biological replicates). All data are shown as mean ± s.e.m.

**Suppl. Figure 2.**
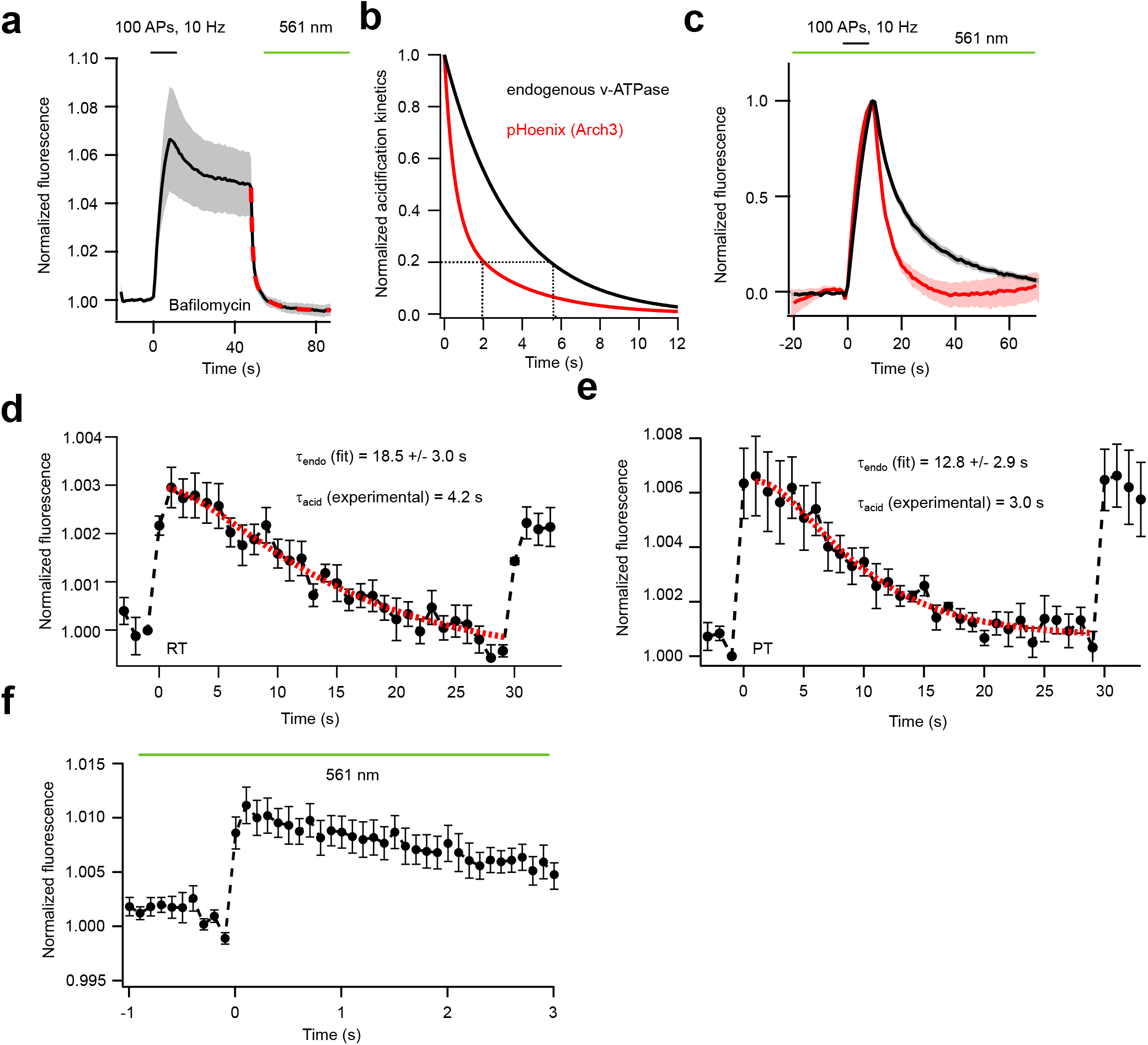
pHoenix characterization and imaging of single vesicle exo-endocytosis with higher sampling rate. **a)** Acidification kinetics of the optogenetic proton pump Arch3 at RT. Hippocampal neurons were stimulated with 100 APs at 10 Hz in presence of Baf. After completion of endocytosis, the proton pump Arch3 was activated by 561 nm laser light to estimate the optogenetically induced acidification rate of pHoenix (n=8 coverslips from 3 biological replicates; red: double exponential fit of the average curve). Fitting the individual curves to a double-exponential function yields τ_fast_ = 1.3 ± 0.2 s (70 ± 4 % of amplitude) and τ_slow_ = 11.2 ± 4.1 s (30 ± 4 % of amplitude). **b)** Comparison of the SV acidification kinetics of the endogenous v-ATPase (black) and light-induced acidification by Arch3 (pHoenix, red) at PT. Shown are the fits of the average traces (Fig 1c and Fig. 2b). Dashed lines indicate time points of 80 % decay. **c)** Average fluorescence response of pHoenix to a stimulus train of 100 APs at 10 Hz at RT (black; n = 7 coverslips from 2 biological replicates) and at PT (red; n = 6 coverslips from 2 biological replicates) with Arch3 activation. **d)** Ensemble-averaged fluorescence traces of neurons expressing pHoenix in response to single APs at RT without light-induced pHoenix activation (n=6 coverslips from 3 biological replicates). A double-exponential fit to the decay curve (red, shown as fit of the average curve) represents SV endocytosis and re-acidification as two sequential first-order processes. τ_acid_ was fixed to 4.2 s during the fitting (experimental results shown in 1d) to obtain τ_endo_. Fitting of the individual curves resulted in τ_endo_ = 18.5 ± 3.0 s. **e)** Ensemble-averaged fluorescence trace of pHoenix responses without light activation to single APs at PT (n=6 coverslips from 3 biological replicates). Fitting the individual curves to the double-exponential model with τ_acid_ fixed to 3 s resulted in τ_endo_ = 12.6 ± 2.9 s. **f)** Ensemble-averaged fluorescence trace of neurons expressing pHoenix at PT with light-induced pHoenix activation in response to single APs sampled at 10 Hz for 4 s around single AP delivery (n=30 coverslips from 6 biological repeats with average of responses to 4 APs at 30 s interval). All data are shown as mean ± s.e.m.

**Suppl. Figure 3.**
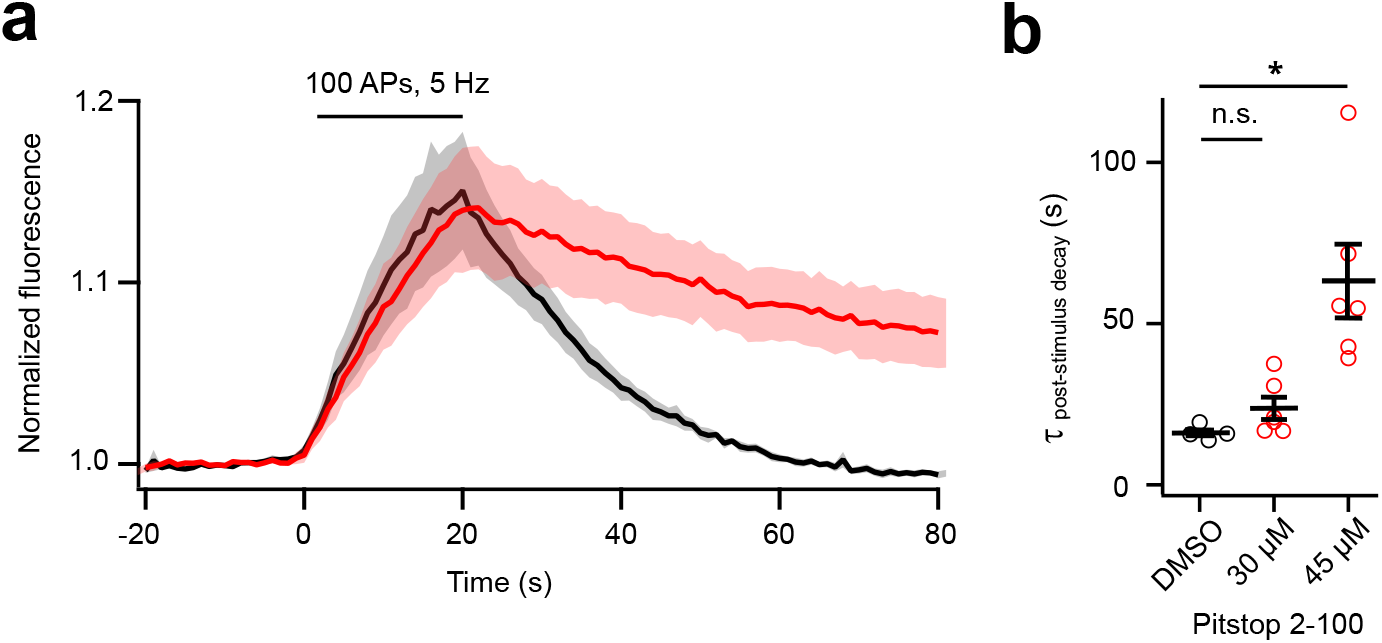
Block of endocytosis by Pitstop 2-100. **a)** Average time course of Syp1-pHl fluorescence in response to 100 APs at 5 Hz for control (DMSO, black) and in presence of 45 µM Pitstop 2-100 (red). **b)** Time constants of fluorescence recovery estimated by mono-exponential fits for different Pitstop 2-100 concentrations (n=5-6 coverslips each from 2 biological replicates). All data are shown as mean ± s.e.m. n.s.: non-significant, * p <0.01, two-tailed unpaired t-test.

## Acknowledgments

We thank K. Tkotz and A. Roetrige for excellent technical support. S.P. was supported by the CIM-IMPRS Graduate School of the University of Muenster, Germany. J.K. was supported by grants of the German Science Foundation (SFB 944 and SFB 1348).

## Author contributions

M.K. and J.K. designed the experiments, S.P. and M.K. conducted the experiments, S.P. and M.K analyzed data, S.P., M.K., and J.K. wrote the paper.

